## Supplementary material for "Structural Insights into Photoactivation of Plant Cryptochrome-2": 082820_SupplInformation.pdf

### Supplementary Information

**Figure S1:** Multiple sequence alignment and conservation analysis of selected cryptochromes and photolyase ortholog.

**Figure S2:** AtCRY2-PHR protein characterization.

**Figure S3:** Structural analysis of CRYs.

**Figure S4:** Characterization of CRYs tryptophan triad and FAD pocket.

**Table S1:** Detailed summary of PHR photo-induced conformational changes.

**Movie S1** and **Movie S2** descriptions.

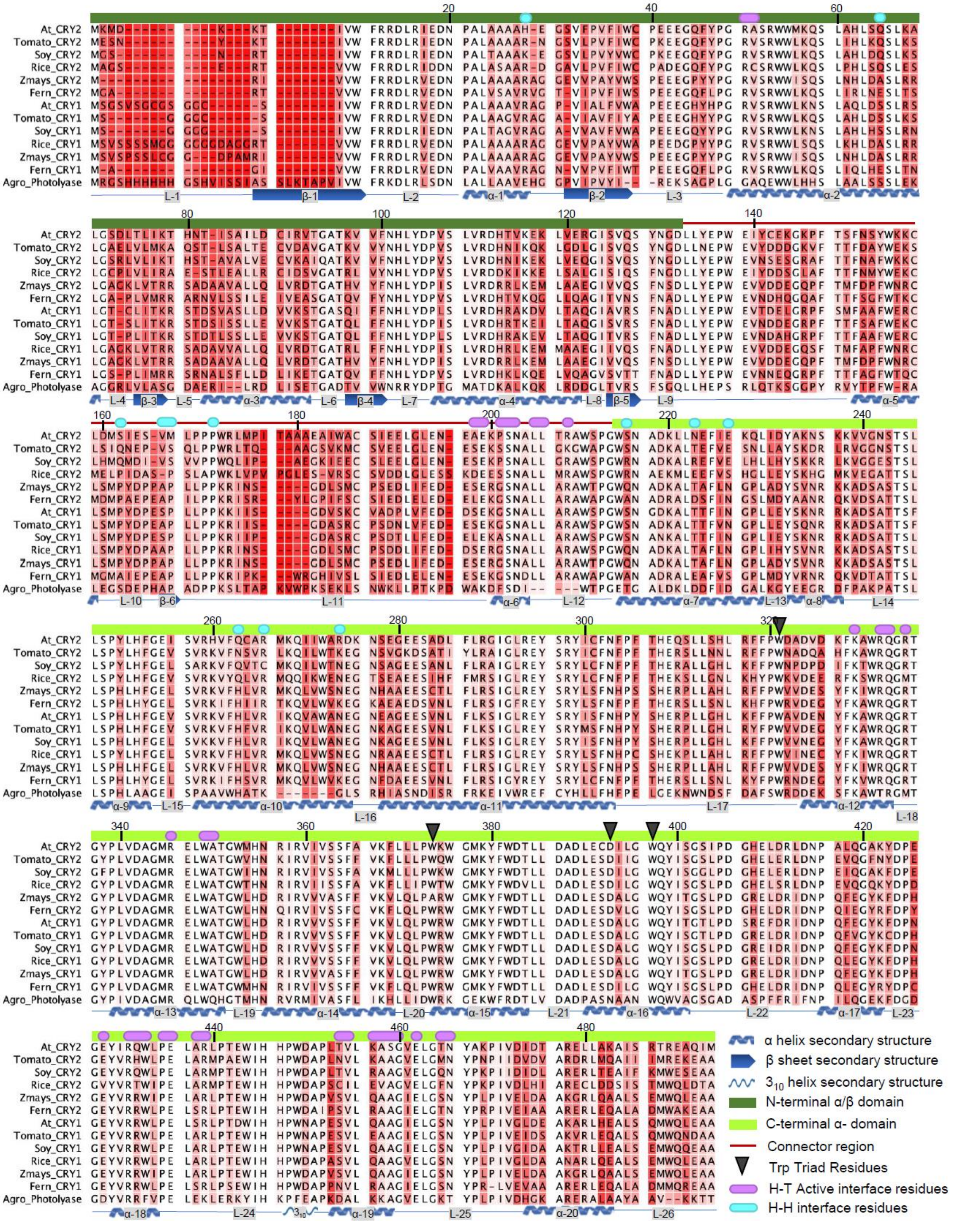

At_CRY2
Tomato_CRY2
Soy_CRY2
Rice_CRY2
Zmays_CRY2
Fern_CRY2
At_CRY1
Tomato_CRY1
Soy_CRY1
Rice_CRY1
Zmays_CRY1
Fern_CRY1
Agro_Photolyase
α helix secondary structure
β sheet secondary structure
3₁₀ helix secondary structure
N-terminal α/β domain
C-terminal α- domain
Connector region
Trp Triad Residues
H-T Active interface residues
H-H interface residues

**Figure S1. Multiple sequence alignment and conservation analysis of selected cryptochromes and photolyase ortholog.** Amino acid alignment of 12 plant cryptochrome proteins with photolyase used as outgroup for comparison. Intensity of red behind residues shows degree of divergence. Numbers on residues refer to position in *At*CRY2-PHR sequence. Key *At*CRY2-PHR subdomains N-terminal α/β domain (dark green), C-terminal α- domain (bright green), and Linker Region (red line) are indicated above alignment. Secondary structure of *At*CRY2-PHR sequence is shown below sequence in blue as alpha helices (α1- α20), beta sheets (β1- β6), $3_{10}$ helices ($3_{10}$) and non-secondary structure-containing loops (L1-L26). Black arrow indicates residues involved in Trp triad. Magenta ovals indicate residues involved in formation of H-T interface and Cyan ovals indicate residues involved in formation of H-H interface. In naming scheme At represents *Arabidopsis thaliana*, Tomato *Solanum lycopersicum*, Soy *Glycine max,* Rice *Oryza sativa*, Fern *Asplenium yunnanense*, Zmays *Zea mays*, and Agro *Agrobacterium tumafaciens*.

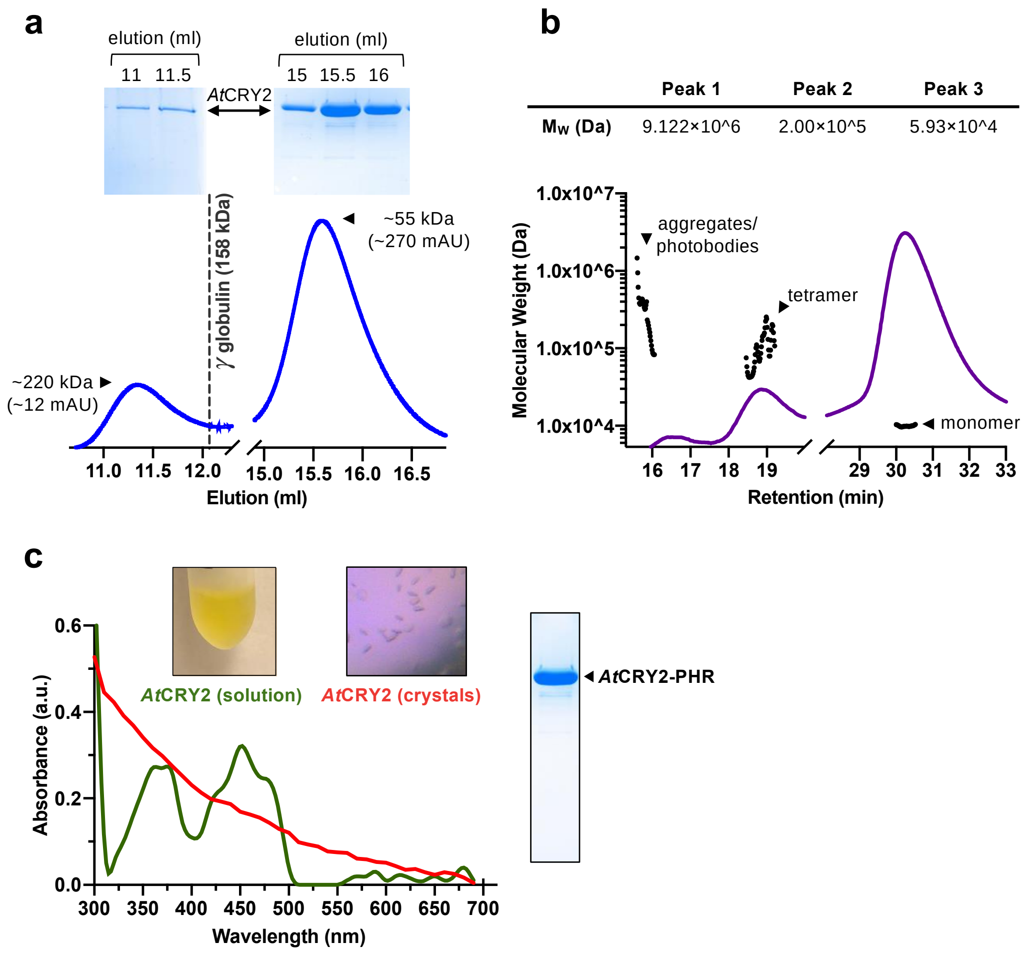


**Figure S2. AtCRY2-PHR protein characterization.** **(a)** Size exclusion analysis of purified *At*CRY2 PHR domain. MW were estimated by biomolecule standard markers. Elution fractions were resolved by SDS-PAGE and Coomassie staining *top* and UV absorption spectra *bottom*. **(b)** SEC-MALS analysis of *At*CRY2-PHR domain. (**c)** Absorption spectra of *At*CRY2-PHR proteins in solution and in crystals was measured at the range of 300 nm to 700 nm wavelength. Sample of the purified *At*CRY2-PHR used for crystallization trials was resolved by SDS-PAGE and visualized via Coomassie staining.

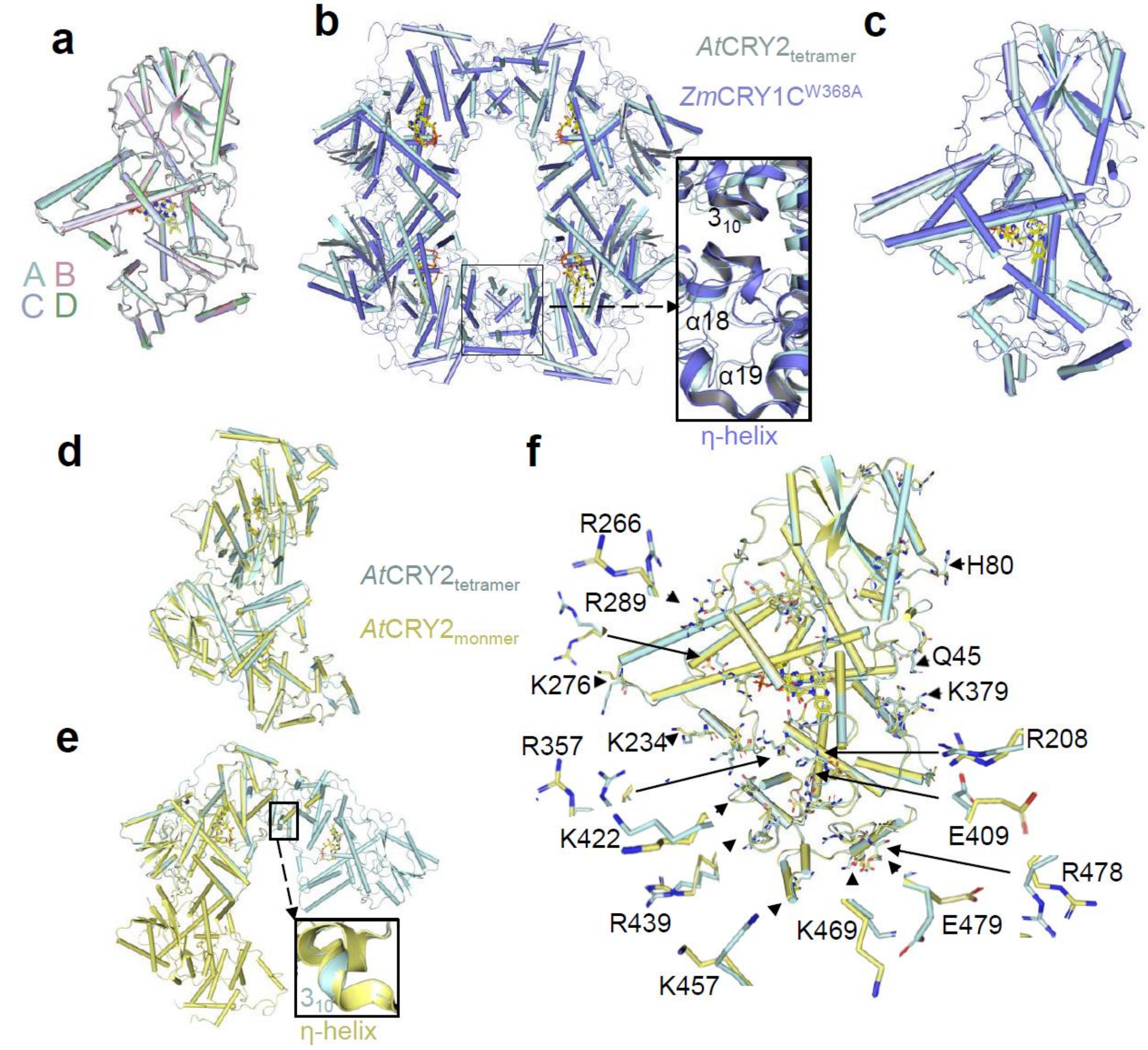


**Figure S3. Structural analysis of CRYs. (a)** Superposition of monomers A (cyan), B (pink), C (purple), D (green) of *At*CRY2-PHR$_{tetramer}$ determined in this study. The r.m.s deviation is calculated for the superposition between monomers A and D (0.59 Å), monomer B and C (0.59 Å) and monomer B and D (0.62Å) chains respectively. **(b-c)** Superposition of *At*CRY2-PHR$_{tetramer}$ (cyan) with mutant *Zm*CRY1C mutant (blue, PDB:6LZ3) resulting in a r.m.s deviation of 1.0Å. Top view of tetramers superposition with close up view on oligomeric interface (b) and side view of monomers **(c)**. **(d-e)** Dimers superposition of *At*CRY2-PHR$_{tetramer}$ and *At*CRY2-PHR$_{monomer}$ (yellow, PDB: 6K8I) with two distinct rearrangements resulting in r.m.s deviation of 0.88 Å for all the Cα atoms (monomer A with D), and r.m.s deviation of 0.86 Å (monomer B with C). **(f)** Superposition of monomers of *At*CRY2-PHR$_{tetramer}$ (cyan) and CRY2-PHR$_{monomer}$ (yellow, PDB: 6K8I). Side chains with distinct structural deviation are labeled and shown as sticks, close ups of key residues are zoomed in and pointed to with black arrows. All CRYs structure represent the PHR domain.

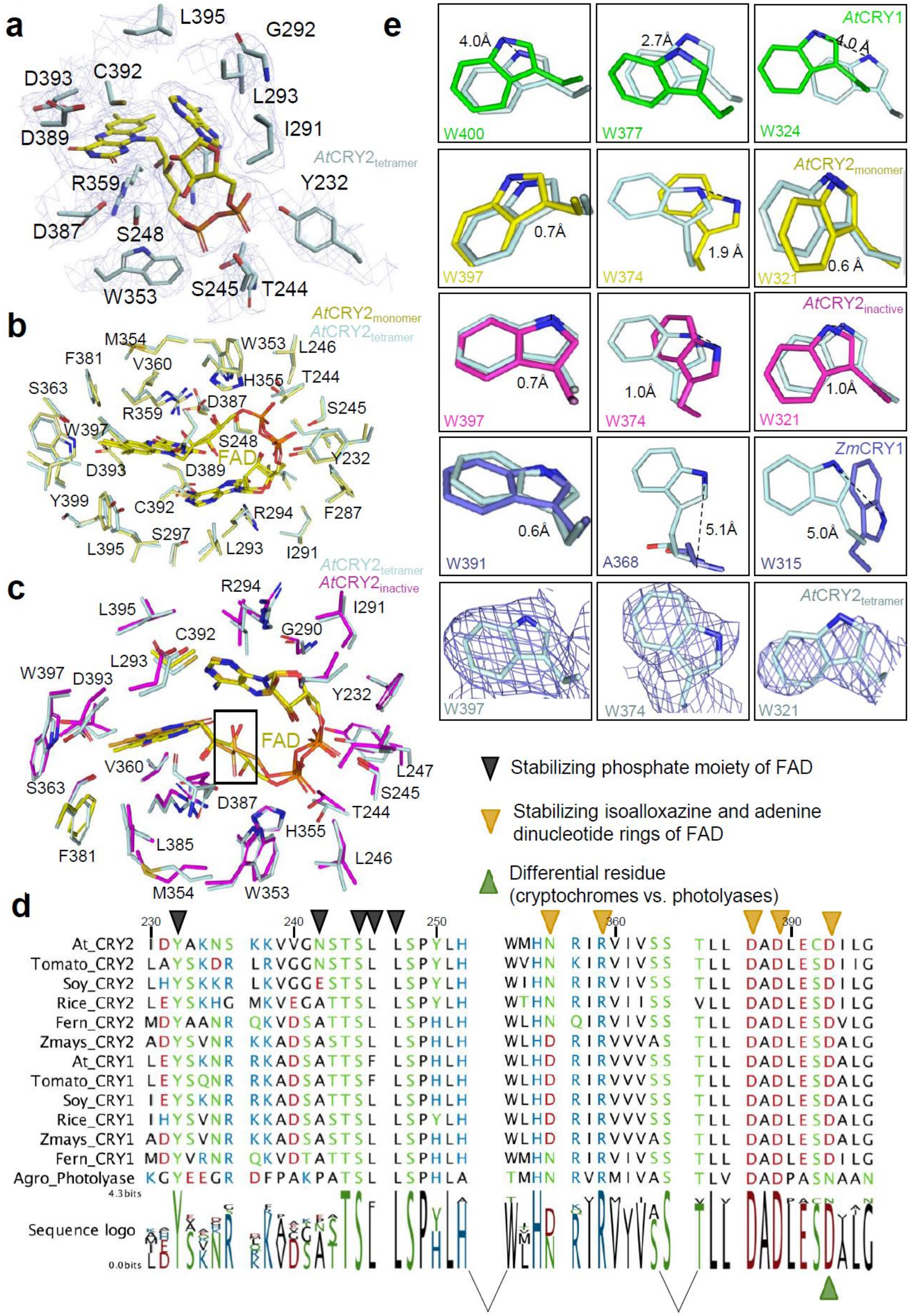
a
L395
G292
D393
C392
L293
D389
I291
AtCRY2tetramer
R359
Y232
D387
S248
W353
S245
T244
b
AtCRY2monomer
AtCRY2tetramer
M354
W353
F381
V360
L246
H355
S363
T244
D387
R359
S245
W397
S248
FAD
D393
D389
Y232
Y399
C392
R294
F287
S297
L395
L293
I291
c
AtCRY2tetramer
AtCRY2inactive
R294
L395
I291
C392
G290
L293
W397
D393
Y232
FAD
V360
L247
S363
D387
S245
T244
H355
L385
F381
L246
M354
W353
d
230
240
250
360
390
At_CRY2 IDYAKNS KKVVGNSTSL LSPYLH WMHN RIRVIVSS TLL DADLECDILG
Tomato_CRY2 LAYSKDR LRVGGNSTSL LSPYLH WVHN KIRVIVSS TLL DADLESDIIG
Soy_CRY2 LHYSKKR LKVGGESTSL LSPYLH WIHN RIRVIVSS TLL DADLESDILG
Rice_CRY2 LEYSKHG MKVEGATTSL LSPYLH WTHN RIRVIISS VLL DADLESDILG
Fern_CRY2 MDYAANR QKVDSATTSL LSPHLH WLHN QIRVIVSS TLL DADLESDVLG
Zmays_CRY2 ADYSVNR KKADSASTSL LSPHLH WLHD RIRVVVAS TLL DADLESDALG
At_CRY1 LEYSKNR RKADSATTSF LSPHLH WLHD RIRVVVSS TLL DADLESDALG
Tomato_CRY1 LEYSQNR RKADSATTSF LSPHLH WLHD RIRVVVSS TLL DADLESDALG
Soy_CRY1 IEYSKNR RKADSATTSL LSPHLH WLHD RIRVVVSS TLL DADLESDALG
Rice_CRY1 IHYSVNR KKADSASTSL LSPYLH WLHD RIRVVVSS TLL DADLESDALG
Zmays_CRY1 ADYSVNR KKADSASTSL LSPHLH WLHD RIRVVVAS TLL DADLESDALG
Fern_CRY1 MDYVRNR QKVDTATTSL LSPHLH WMHN RIRVIVSS TLL DADLESDILG
Agro_Photolyase KGYEEGR DFPAKPATSL LSPHLA TMHN RVRMIVAS TLV DADPASNAAN
4.3bits
Sequence logo
0.0bits
Stabilizing phosphate moiety of FAD
Stabilizing isoalloxazine and adenine dinucleotide rings of FAD
Differential residue (cryptochromes vs. photolyases)
e
AtCRY1
4.0Å
2.7Å
4.0 Å
W400
W377
W324
AtCRY2monomer
0.7Å
1.9 Å
0.6 Å
W397
W374
W321
AtCRY2inactive
0.7Å
1.0Å
1.0Å
W397
W374
W321
ZmCRY1
0.6Å
5.1Å
5.0Å
W391
A368
W315
AtCRY2tetramer
W397
W374
W321

**Figure S4. Characterization of CRYs tryptophan triad and FAD pocket. (a)** 2Fo-Fc electron density map (1σ cut-off) (mesh light blue) of FAD (yellow) binding pocket within *At*CRY2-PHR$_{tetramer}$ (cyan) with the direct interacting amino acids residues shown as sticks. (**b**) Close up view on the superposition of *At*CRY2-PHR$_{tetramer}$ with *At*CRY2-PHR$_{monomer}$ (PDB: 6K8I) within the FAD binding pocket. r.m.s deviation of 0.4 Å. (**c**) Close up view on the superposition of *At*CRY2-PHR$_{tetramer}$ with *At*CRY2-PHR$_{inactive}$ (PDB: 6K8K) within the FAD binding pocket. r.m.s deviation of 0.4 Å. The box highlighted in black shows the flip in the atoms O3 and O4 of FAD (orange) with respect to *At*CRY2-PHR$_{tetramer.}$ (**d**) Sequence alignment and conservation of cryptochrome FAD binding pocket across plant species. Colors indicate amino acid polarity. **(e)** Structural analysis of *At*CRY2-PHR$_{tetramer}$ (light blue) tryptophan triad superposition with *At*CRY1 (green, PDB: 1U3C), *At*CRY2-PHR$_{monomer}$ (yellow, PDB: 6K8I), *At*CRY2-PHR$_{inactive}$ (Magenta, PDB:6K8K), *Zm*CRY1C mutant (Blue, PDB:6LZ3). All CRYs structure represent the PHR domain.

### Table S1. Detailed summary of PHR photo-induced conformational changes

| Major structural changes | | Residue positions | Proposed Function | References |
|---|---|---|---|---|
| Presence of a short helix | | 189-192 (*At*CRY2-PHR$_{inactive}$)*<br>181-184 (*At*CRY2-PHR$_{inactive}$) | Photo-inactivation | This study and [16,32,48] |
| | | 189-192 (*At*CRY2-PHR$_{monomer}$)** | | |
| | | 187-190 (*At*CRY1-PHR) | | |
| | | 144-146 (*At*CRY1-PHR) | | |
| Absence of a short helix | | 189-192 (*At*CRY2-PHR$_{tetramer}$)<br>181-184 (*At*CRY2-PHR$_{tetramer}$) | Photo-activation and oligomerization | This study and [16,32,48] |
| Presence of 3$_{10}$ helix | | 448-450 (*At*CRY2-PHR$_{tetramer}$) | Photo-activation and oligomerization | This study |
| α19 helix | Change of 0.32Å deviation | 453-459 (*At*CRY2-PHR$_{inactive}$) | Participate in oligomeric interface | This study [16,32,48] |
| | Change of 0.41Å deviation | 453-459 (*At*CRY2-PHR$_{monomer}$) | | |
| | Change of 0.39 deviation | 455-463 (*At*CRY1-PHR) | | |
| α18 helix | Change of 0.32Å deviation | 427-433 (*At*CRY2-PHR$_{inactive}$) | Participate in oligomeric interface | This study [16,32,48] |
| | Change of 0.32Å deviation | 427-433 (*At*CRY2-PHR$_{monomer}$) | | |
| | Change of 0.32Å deviation | 430-437 (*At*CRY1-PHR) | | |
| α13 helix | Change of 0.20Å deviation | 340-350 (*At*CRY2-PHR$_{inactive}$) | Participate in oligomeric interface | This study [16,32,48] |
| | Change of 0.19Å deviation | 340-350 (*At*CRY2-PHR$_{monomer}$) | | |
| | Change of 0.21Å deviation | 342-354 (*At*CRY1-PHR) | | |
| α12 helix | Change of 0.24Å deviation | 326-333 (*At*CRY2-PHR$_{inactive}$) | Participate in oligomeric interface | This study [16,32,48] |
| | Change of 0.20Å deviation | 326-333 (*At*CRY2-PHR$_{monomer}$) | | |
| | Change of 0.15Å deviation | 328-337 (*At*CRY1-PHR) | | |
| α6 helix | Change of 0.4Å deviation | 201-209 (*At*CRY2-PHR$_{inactive}$) | Participate in oligomeric interface | This study [16,27,32,48] |
| | Change of 1.3Å deviation | 201-299 (*At*CRY2-PHR$_{monomer}$) | | |
| | Change of 1.0Å deviation | 199-213 (*At*CRY1-PHR) | | |

* *At*CRY2-PHR$_{inactive}$ represents the molecular structure of CRY2-PHR in complex with BIC2

** *At*CRY2-PHR$_{monomer}$ represents the molecular structure of CRY2-PHR in monomeric form

### Movie S1. Structural movements between active an inactive CRY2

Morph movie of the conformational variation of the interconnecting loop between active and inactive CRY2 (PDB: 6K8K).

### Movie S2. Overall structural variations between active an inactive CRY2

Morph movie showing the overall structural variation between the active (pale cyan) and inactive CRY2 (PDB: 6K8K) structures. *Left screen:* overall structural variations based on superposition. *Right screen:* close up view on amino acids residues involved in electron transport pathway are represented in sticks.
